# Targeted Memory Reactivation of Face-Name Learning Depends on Ample and Undisturbed Slow-Wave Sleep

**DOI:** 10.1101/2021.02.16.431530

**Authors:** Nathan Whitmore, Adrianna M. Bassard, Ken A. Paller

## Abstract

Face memory, including the ability to recall the name of a familiar person, is often crucial in social interactions, and like many other memory functions, it may rely on sleep. We investigated whether targeted memory reactivation during sleep could improve associative and perceptual aspects of face memory. Participants studied 80 face-name pairs, and then a subset of spoken names was presented unobtrusively during a daytime nap. This reactivation preferentially improved recall for those face-name pairs, as modulated by two factors related to sleep quality. That is, the memory benefit was positively correlated with the duration of stage N3 sleep (slow-wave sleep) and with the extent to which cues presented during SWS did not produce a sleep disruption indexed by increased alpha-band electroencephalographic activity in the 5 seconds after a cue. Follow-up analyses showed that a memory benefit from presenting spoken names during sleep was evident in participants with high amounts of SWS or with low amounts of sleep disruption. We conclude that sleep reactivation can strengthen memory for specific face-name associations and that the effectiveness of reactivation depends on uninterrupted N3 sleep.

## Introduction

Successful face recognition and name recall — such as when you see your friends in a crowd and call to them by their names — are important in social contexts. People are extraordinarily adept at recognizing faces of individuals they’ve met even just once. There are also times when people fail to recognize someone they know quite well. And it can be exceedingly embarrassing to forget a name that should have been remembered.

What determines which memories continue to be enduringly available and which are forgotten? Given that the human brain is remarkably active during sleep, researchers have repeatedly asserted that neural events during sleep may function to stabilize and strengthen recently acquired memories (e.g., Marr, 1971; Winson, 1984; Paller, 1997; Paller, Mayes, Antony, & Norman, 2020). The delineation of these neural events and their specific ramifications for memory has become increasingly central to memory research and the science of learning.

A prevalent view is that memories can benefit due to spontaneous replay during sleep (e.g., Born and Wilhelm, 2012; Sirota et al., 2003; Sejnowski and Destexhe, 2000; Pavlides and Winson, 1989). In recent years, Targeted Memory Reactivation (TMR) has emerged as a useful tool for investigating this process (Oudiette and Paller, 2013). In the TMR procedure, information that people learn is associated with a sound or smell during learning. Researchers then present the same sensory cue while people sleep, without waking them. After sleep, people remember information associated with the cue stimulus better than non-associated information, a consistent finding confirmed in a recent meta-analysis (Hu et al., in press). Moreover, the notion that TMR benefits memory through reactivation is supported by neuronal evidence of hippocampal place cell replay engaged following the presentation of learning-related sounds during sleep (Bendor and Wilson, 2012).

TMR offers the potential to enhance memory with a simple, non-invasive intervention during sleep, which may be useful in many scenarios. Because learning face-name associations is an important and widely relevant form of memory, we asked whether TMR could enhance this type of learning.

We developed a procedure whereby participants learned about people ostensibly in either a Japanese-history class or a Latin-American-history class. Learning was accompanied by a background music track, either traditional Japanese music or traditional Latin-American music, respectively. Each classroom had 40 pupils. To learn the names of these pupils, participants viewed each face adjacent to the corresponding written name while also hearing the spoken name. Memory testing, feedback, and visualization practice served to solidify learning. We assessed face recognition as well as name recall. Then, after a period of sleep during which some of the spoken names were softly presented, we assessed memory again. Figure 1 shows the experimental procedure schematically, and additional details are provided in *Materials and Methods*.

**Figure 1.**
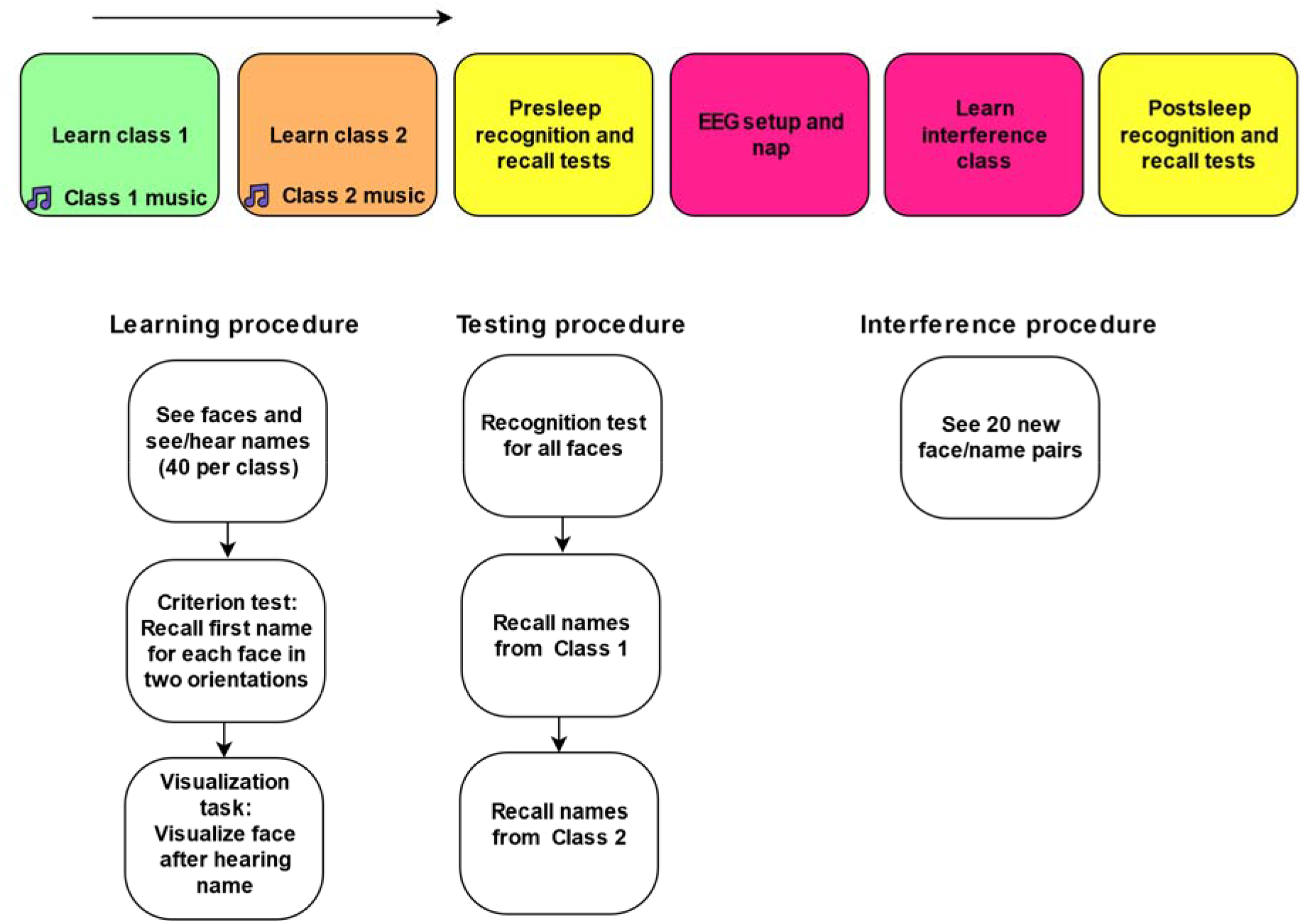
Experimental paradigm overview.

Our results showed that the ability to recall the correct name when viewing the corresponding face was improved when that memory was reactivated during sleep, and that this memory improvement was modulated by the integrity and duration of slow-wave sleep. The findings suggest that effective memory reactivation with these procedures requires sufficient slow-wave sleep not interrupted by alpha activity in the first few seconds after cue presentation.

## Results

### The influence of TMR on recall varied with duration of stage N3 sleep

On the name recall test, participants correctly recalled a mean of 74.25 names before sleep and 75.00 names after sleep (in each case from a total of 80 face-name associations). When requested, hints were provided in the form of starting letters, but participants were correct with no hints for 47.00 names before sleep and for 46.71 names after sleep. Across all 80 pairs in the test, participants requested a mean of 0.76 hints per pair before sleep and 0.80 after sleep. Overall, recall performance was very accurate, and very similar before and after sleep.

To measure small changes in recall performance, we developed a sensitive measure of recall, called *alternatives per correct* (APC). Unlike a binary measure of whether name recall was simply correct or incorrect, APC provides a graded measure of performance for each name by considering how much information the participant requests before making a correct response. APC is computed as the number of names not yet ruled out from the initial pool of 80 names based on the hints received. For example, if the correct name was recalled after receiving the first letter, *S*, and there were 8 names in the pool starting with *S*, then the APC would be 8. If the correct name was recalled after receiving two letters, *ST*, and there were 3 names in the pool starting with *ST*, then the APC would be 3. Thus, a high APC indicates superior recall compared to a low APC. APC was set to 0 if recall was not correct and was computed only for trials in which hints were given (39% of all trials).

To ascertain whether sleep influenced name recall when hints were required, we relied on the change in APC score across sleep (ΔAPC). The mean APC was 2.18 pre-sleep and 2.35 postsleep. ΔAPC did not differ significantly between the cued class, designated *class C*, and the uncued class, designated *class U* [mean ΔAPC 0.40 and −0.06, respectively; *t*(23)=1.48, *P*=0.15]. However, the size of the *ΔAPC cueing effect* (difference between class-C ΔAPC and class-U ΔAPC) was highly correlated with the duration of stage N3 sleep [*r*(22)=0.56, P=0.004]. As shown in Figure 2, this correlation was consistent with the results from a median split on stage N3 duration, which revealed a significant cueing effect for the high-N3 participants [*t*(11)=2.83, *P*=0.016], but not for the low-N3 participants [*t*(11)=1.06, *P*=0.31]. High-N3 participants had a mean of 43 min of N3 whereas low-N3 participants had a mean of 19 min of N3.

**Figure 2.**
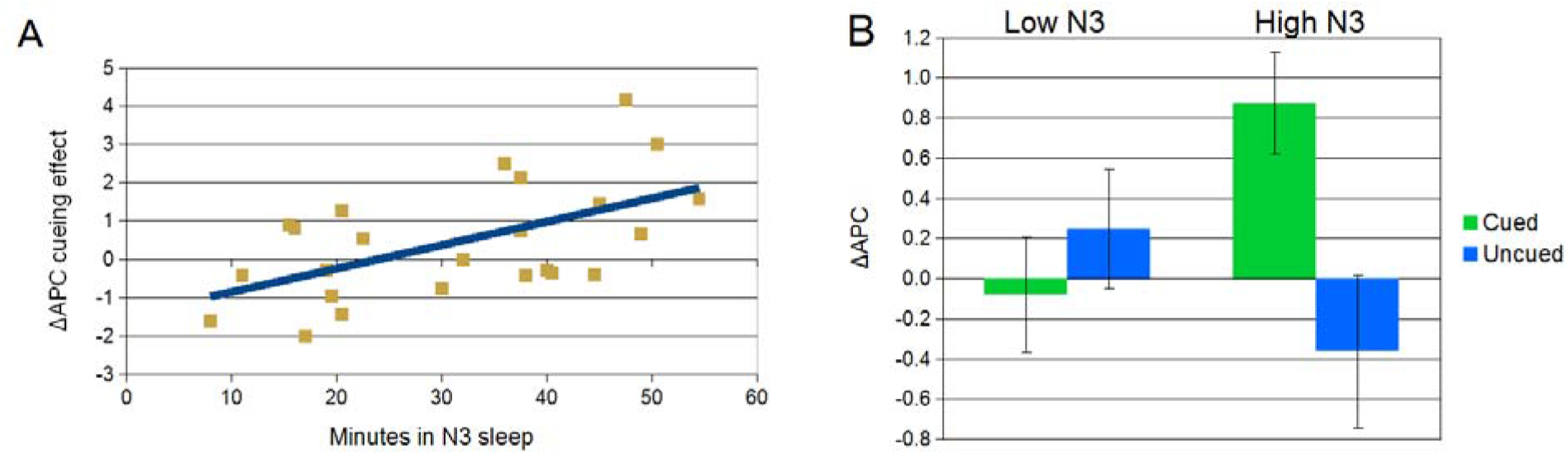
(A) N3 duration was linearly correlated with ΔAPC cueing effect for the cued class relative to the uncued class. (B) Effects of TMR on recall were evaluated via a median split of participants by duration of stage N3 sleep. Recall as assessed by ΔAPC differed as a function of cueing in participants with high N3 duration but not those with low N3 duration.

We also investigated whether TMR improved the total number of names recalled. Neither the number of names participants recalled nor the number of names recalled with no hints differed as a function of cueing [*t*(23)=0.93,0.88, *P*=0.36,0.39 respectively]. The cueing effect on total names recalled and names recalled without hints did not correlate with stage N3 duration [*r*(22)=0.24,0.2, *P*=0.3, 0.35 respectively].

The design also enabled comparisons between cued and uncued names within class C. However, we did not observe differences between cued/uncued names using either APC or number of names recalled [*t*(15)=0.21, *P*=0.83, and *t*(15)=1.1973, *P*=0.25, respectively; for both tests data were available for this analysis from 16 participants due to an issue with coding the cued and uncued items]. In both cases, differences between cued and uncued names did not correlate with stage N3 duration.

### The influence of TMR on recall depended on undisturbed sleep

A previous report found that participants who self-reported sleep disruption during TMR did not benefit from memory cues (Göldi & Rasch, 2019). To objectively assess sleep disruption related to cue delivery during sleep, we quantified arousals in the EEG after spoken name cues. As an arousal is conventionally defined by a sharp shift in the spectral content of the EEG (Iber et al, 2007), we summed the absolute change in EEG power at Cz (increase or decrease) across the frequency band from 0.38-20.35 Hz during the 5 seconds after cue onset relative to the 5 prior seconds. We then used a correlational analysis to examine whether the memory benefit of TMR was associated with this index of sleep disruption.

As shown in Figure 3, cue-related sleep disruption was negatively correlated with the ΔAPC cueing effect [*r*(22)=0.55, *P*=0.006]. Seven participants had a markedly high disruption index [mean disruption=0.23 for these high-disruption participants, 0.11 for all others]; these participants exhibited a nonsignificant ΔAPC cueing effect [mean=-0.63, *t(*6)=1.53, *P*=0.18]; participants with lower disruption exhibited a significant ΔAPC cueing effect [mean=0.9, t(16)=2.57,*P*=0.02].

**Figure 3:**
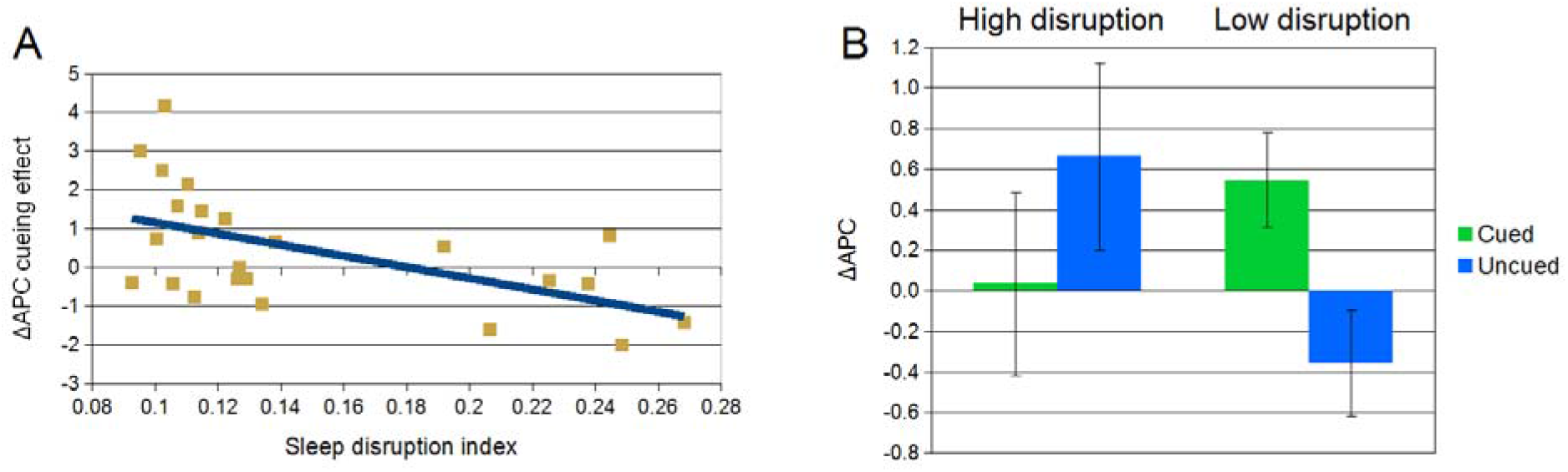
Sleep disruption was inversely correlated with ΔAPC cueing effects for the cued class relative to the uncued class. Sleep disruption formed a bimodal distribution, with a subgroup of markedly disturbed participants (*n*=7). (B) Only the low-disruption subgroup showed improved recall performance for cued items.

Disruption was also negatively correlated with time in N3 [*r*(22)=0.6, *P*=0.002]. Multiple regression revealed that N3 duration and disruption index were redundant predictors of the ΔAPC cueing effect, with neither significantly predicting the ΔAPC cueing effect when controlling for the other [*t*(21)=1.7, −1.54; *P*=0.1, 0.14, respectively].

To identify the specific frequencies driving this correlation, we divided the 0.38-20.35 Hz spectrum into six frequency bands, each 3.33-Hz wide, and correlated cue-evoked power in each band with the ΔAPC cueing effect. Low alpha power (7.04-10.37 Hz) was significantly correlated with the cueing effect; a greater cueing effect was seen with less alpha after a cue [*r*(22)=0.48, *P*=0.02]. Cueing effects on ΔAPC were not correlated with changes in any of the other five frequency bands.

### TMR did not affect recognition performance

We investigated whether TMR affected the ability to discriminate previously seen faces from unseen ones (recognition accuracy), to assign previously seen faces to the correct class (source recognition accuracy), and to discriminate previously seen faces in a new orientation. Because of technical shortcomings, usable recognition data were available from only 21 participants.

Prior to sleep, participants correctly recognized a mean of 151.3 (95%) of the old faces, assigned 113.6 (71%) to the correct class (Latin or Japanese), and endorsed 51.2 (73%) of the new faces as new. After sleep, participants recognized 150 (94%) of the old faces, assigned 119 (74%) to the correct class, and correctly endorsed 60.6 (87%) of the new faces.

Recognition cueing effects were calculated by comparing the change across sleep for class C (ΔC) to the change across sleep for class U (ΔU) for old faces presented in a previously learned orientation. There was no significant cueing effect on recognition accuracy [ΔC=-0.29, ΔU=-1.05 faces endorsed, *t*(20)=0.89, *P*=0.38] or source recognition accuracy [ΔC=4.1, ΔU=3.52 faces assigned to correct class, *t*(20)=0.23, *P*=0.82]. Results for rotated faces are described in the next section.

We also tested whether cueing effects on recognition depended on N3 duration and quality. Neither recognition accuracy nor source recognition accuracy correlated with N3 duration [*r*(19)=0.0,0.003, *P*=1,0.99 for recognition and source recognition respectively] or sleep disruption index [*r*(19)=0.04,0.19, *P*=0.85,0.41 for recognition and source recognition respectively].

### TMR did not improve recognition of rotated faces

Given previous results that sleep can improve implicit aspects of face memory (Wagner et al, 2003), we tested whether cues improved the implicit learning of face structure by testing participants ability to recognize previously viewed faces in a new orientation (rotated faces). Reaction times for rotated faces declined slightly, but not significantly, across sleep [mean = 1.71 s pre-sleep, 1.63 s post-sleep, *t*(20)=1.42, *P*=0.17). Also, the change in rotated-face recognition time did not differ as a function of cueing [*t*(20)=0.05, *P*=0.96], and the cueing effect did not correlate with SWS duration [*r*(19)=0.08, *P*=0.74] or sleep disruption [*r*(19)=0.03, *P*=0.9].

We also considered whether TMR might selectively improve the accuracy of rotated face recognition. Participants did not endorse more rotated faces from class C than from class U [*t*(20)=1.4, *P*=0.18] and there was no correlation between the recognition cueing effect and N3 time [*r*(19)=0.0, *P*=0.98] or sleep disruption index [*r*(19)=0.13, *P*=0.57].

## Discussion

After a short period of training with arbitrary face-name associations, participants in this experiment were highly successful in recalling the correct name for each face, albeit with the use of letter hints that were available upon request. With these hints, participants recalled 93% of the names. To quantify name-recall success for trials when recall required hints, we relied on APC, a highly sensitive index of the extent to which letter hints helped participants narrow down the number of alternative names. Analyses of this APC measure indicated that when slow-wave sleep was ample and undisturbed, delivering name cues during sleep improved the ability to recall those names when prompted with the corresponding faces.

Delivering auditory or olfactory cues during slow-wave sleep with the goal of targeted memory reactivation has previously been shown to improve many types of memory (Hu et al., in press). In this research area, learning of face-name associations has not previously been examined. Our results add to this literature by showing that an additional type of memory, name recall, is subject to memory reactivation during sleep. More importantly, the experiment also provided novel evidence that might be generally applicable with respect to the relevance of memory reactivation during sleep to any type of learning. That is, objective measures of N3 duration and sleep disruption after cues moderated the degree of name-recall improvement from TMR.

Compatible findings have been reported from studies of TMR in word-learning paradigms using self-reports of sleep disruption (Göldi & Rasch, 2019) and using REM duration (Batterink, Westerberg, & Paller, 2017). Furthermore, N3 duration in the present experiment had a strong negative correlation with arousal after cues, suggesting that N3 duration and phasic sleep disruption may index a common factor. We therefore propose that variation in these two aspects of SWS quality represent factors sometimes overlooked in the broader TMR literature. Such variation may explain some instances when TMR does not benefit memory.

Our results raise a further question: what is the relationship between momentary arousal, N3 duration, and TMR effects? Göldi and Rasch (2019) proposed that sleep disruption caused by auditory stimulation may introduce interference into newly reactivated memories. Conversely, arousals may have no direct effect on memory, but simply reduce N3 duration, which in turn reduces TMR benefits. A third possibility is that short N3 duration and high arousability both reflect a latent factor of sleep quality that mediates TMR effects. A future experiment that included a direct manipulation of arousal could be used to more directly test these possibilities.

In this experiment, TMR improved memory only on the APC measure, which was developed with the intention of quantifying small memory changes with high resolution. We did not observe improvements for accuracy in other measures of recall, suggesting that the selective enhancement of the cued class was relatively small. We also did not observe effects on our recognition tests, consistent with meta-analytic results showing that TMR does not have a reliable effect on recognition measures (Hu et al., in press). The effects on recall tests may have been limited by several factors, including ceiling effects for the total number of items recalled, a limited opportunity for forgetting to occur between the two memory tests, and unintended spreading of reactivation from the cued class to the uncued class. Modified protocols, for instance with memory testing at a longer delay or utilizing a between-subjects design, may shed light on the role of these factors. When memory is tested immediately after sleep with TMR, evidence that reactivation was helpful might be most clearly evident for memories that fall just short of being strong enough to be remembered at that time point.

In our procedure, sleep cues included both specific names from the learning phase as well as the background track for half of the name learning, which was a unique music genre. We therefore cannot determine whether TMR benefits were due to the spoken names, the music, or both. In prior experiments, TMR benefits have been observed using spoken words (e.g., Schreiner & Rasch, 2015; Cairney et al., 2018) as well as short music tracks (e.g., Antony et al., 2012; Sanders et al., 2019). Interestingly, we did not observe differences in TMR effects between the cued and uncued items within the cued class. This outcome suggests that reinstating the music track played during learning reactivated the corresponding class during sleep, improving memory for both cued and uncued names within that class. Similar results have been observed in TMR studies using spatial tasks with contextual odor cues (e.g., Rasch et al., 2007). Alternatively, we may have failed to detect within-class TMR effects due to insufficient power or an insufficient number of trials in the cued and uncued conditions.

Overall, the present results show that provoking memory reactivation during sleep can influence the learning of face-name associations, with consequences for whether people can recall the correct name after sleep. Furthermore, the magnitude of this effect was found to depend on the length of N3 sleep and on the absence of arousal after cues. We propose that N3 duration and cue-evoked arousal are important factors shaping how memory reactivation during sleep influences subsequent memory performance. These factors thus have relevance for future research aimed at making progress in understanding the neural mechanisms whereby learning is influenced by subsequent sleep even when there is no sensory input during sleep. Considerations of N3 disruption might be especially relevant for circumstances when memory modification is desirable, including clinical applications (e.g., Paller, 2017), wherein further studies are needed to determine if methods of targeted memory reactivation could prove helpful.

## Materials and Methods

### Participants and Experimental Design

Participants (*N*=24) were 8 males and 16 females 18-31 years old. Figure 1 shows an overview of the procedure, which was approved by the university institutional review board. The design provided within-subject comparisons as a function of whether cues were presented during sleep. After participants arrived in the lab and gave written informed consent, the following phases transpired: learning, pre-nap test, bioelectric recording setup, nap, and post-nap test.

### Procedure

#### Learning Phase

Participants completed a face-name learning task intended to simulate learning the names of pupils in two classes, with 40 face-name pairs per class. Each class was associated with a distinct instrumental music track, traditional Latin American music or traditional Japanese music (see Supplementary Materials). Participants completed all parts of the learning tasks for Class 1 followed by all parts of the learning tasks for Class 2.

Face-name learning started with initial exposure plus interspersed name recall. Participants viewed sets of five face-name pairs, each shown one at a time while the spoken name was played over speakers. In addition to the name, which included a first name and a surname, a one-sentence fact about the person (such as “I love cats and fall weather”) was also viewed. Each to-be-learned person was presented twice during initial exposure with the same biographical information, with the face was shown once in a quarter-right and once in a quarter-left orientation, to maximize learning of the 3D structure. Prompts for name recall appeared after every set of five initial exposures. While prompted with a face, participants attempted to recall the corresponding name, with feedback following each recall attempt.

Following initial exposure with interspersed name recall, participants trained to criterion in the criterion test with all 40 faces. Training was complete when the participant gave the correct first name for each person in the class in both orientations.

Participants were then asked to visualize each person in the class when the corresponding name was spoken. This part of the procedure was intended to promote face visualization in response to names presented during sleep. Each visualization attempt was followed by a prompt to identify the visualized person from two same-sex alternatives. The two faces were presented 3.5 s after name onset. Feedback was presented after the participant made each choice. Following training for Class 1, all training steps were repeated for Class 2, with the order of the Japanese-history class and the Latin-American-history class counterbalancing across subjects. Participants completed all learning steps for both classes in approximately 45 minutes.

#### Pre-sleep and post-sleep tests

After the face-name learning task, a self-paced recognition test with all 80 faces was given. Participants viewed 230 faces sequentially and were asked to decide if each person was in Class 1, Class 2, or was a new person not seen before. The test included 80 previously learned faces (*old*), 70 new faces (*new*), and 80 previously learned faces from a different angle (*rotated*). Old faces were presented at quarter-right orientation, whereas rotated faces were profile views, facing right in the pre-sleep test and left in the post-sleep test. In each test, new faces were presented with equal numbers at quarter-right orientation and the rotated orientation (profile left or profile right) for that test.

Name recall followed recognition testing. Participants were asked to type a person’s first name given a picture of the corresponding face. Faces appeared in either a quarter-right or quarter-left orientation one at a time in a random order. Participants could receive hints by pressing the tab button to receive a letter, up to the first three letters of the person’s first name. No feedback was provided during this test. Each participant performed a cued recall test for Class 1 and then for Class 2.

The same recognition and recall tests were also given starting approximately 10 min after the end of the sleep phase, immediately following an interference-induction task. In this task, participants were asked to memorize 20 new face name/pairs in a simplified procedure with no recall step or biographical details. This manipulation was intended to increase difficulty and simulate the type of interference that occurs in real-world face-name learning.

#### Physiological recording

We recorded electroencephalogram (EEG), electrooculogram (EOG), and electromyogram (EMG) signals during the nap using a BioSemi Active2 system with 32 scalp channels and 4 electrodes on the face. Data were acquired with a sampling rate of 512 Hz, filtered between 0.1 and 100 Hz during recording, and re-referenced to the right mastoid. EOG was recorded with electrodes lateral to the left and right outer canthus and underneath the right eye. EMG was recorded from the chin. Setup and recording began immediately before the start of the sleep period and electrodes were removed after sleep and before the interference task.

#### Nap and TMR

After EEG setup, participants slept on a futon in the same chamber where they completed the learning and testing tasks, with background white noise at a low level (43-44 dB). Participants slept for a mean of 59 min (range: 32 to 92 min). TMR began after participants had been asleep for a mean of 7.3 minutes (range: 1.6 to 21 minutes). TMR was manually initiated when the experimenter visually detected signs of stage N3, which has characteristic EEG slow waves. TMR was paused when sleep transitioned to any other sleep stage or to wake. Offline, raters blind to when cueing occurred determined sleep stages according to standard rules for adults (Iber et al, 2007).

The specific TMR cues presented were selected as follows. Class 1 was randomly assigned to be cued for half of the subjects and Class 2 for the other half. There was thus a cued class and an uncued class (*Class C* and *Class U*, respectively). During TMR, the background music presented while learning class C was played continuously at low volume and half of the spoken names from Class C were presented at 10-s intervals. The specific names played during sleep were chosen to match presleep recall accuracy and number of hints required between cued and uncued names in class C. Intensity of the spoken names was controlled manually to deliver cues at the highest volume possible without causing arousal (45-50 dB peak).

### Behavioral data analysis

We first tested whether cueing differentially influenced memory by comparing the change in memory performance across sleep for class C to that for class U. We defined the primary *cueing effect* by taking the difference between the two change scores as follows: (class C postsleep – class C presleep) – (class U postsleep – class U presleep). We also tested whether cueing differentially influenced memory for cued names in class C versus uncued names in class C.

We computed the number of correctly recognized faces in the recognition test and the number of names correctly recalled in the recall test. We developed a graded measure of recall—the *alternatives per correct* or APC score—to measure small changes in recall ability. The APC score was designed to account for the number of options ruled out by hints before a correct answer was produced. If the participant failed to correctly recall a name, the APC would be zero, and APC would be undefined if the participant recalled the correct name without any hints.

### Arousal analysis

Arousal during sleep is conventionally defined as an abrupt shift in the spectrum of the EEG (Iber et al, 2007). Therefore, we quantified arousal after TMR cues by measuring the absolute difference of the power spectra in two windows: [-5 0] seconds relative to cue onset and [0 5] seconds relative to cue onset. Arousal was calculated as the mean of abs(1-(post cue power/pre cue power)) across linear-spaced 0.256-Hz-wide frequency bins from 0.38 to 20.35 Hz. Power was calculated using a short-time FFT (newtimef, EEGLAB 14.1.1b, 2-s window). As a secondary analysis to identify the frequencies correlated with TMR effects, we divided this frequency range into six 3.33-Hz-wide bins and correlated the event-related spectral-power change for each bin (postcue power / precue power) with the cueing effect.

## Acknowledgements

We gratefully acknowledge grant support from the Mind Science Foundation, NSF BCS-1921678, and NIH T32-MH06756.

## Data Availability Statement

The data that support the findings of this study are available from the corresponding author upon reasonable request.

## Author Contributions

N.W. and K.A.P. designed the experiment. N.W. and A.M.B. collected, analyzed, and interpreted data. K.A.P. supervised the project. N.W. and K.A.P. wrote the manuscript with input and feedback from all authors.

## Competing Interests

The authors declare that there are no competing interests.

## Notes

### Competing Interest Statement

The authors have declared no competing interest.

